# dupRadar: a Bioconductor package for the assessment of PCR artifacts in RNA-Seq data

**DOI:** 10.1101/046243

**Authors:** Sergi Sayols, Denise Scherzinger, Holger Klein

## Abstract

**Background:** PCR clonal artefacts originating from NGS library preparation can affect both genomic as well as RNA-Seq applications when protocols are pushed to their limits. In RNA-Seq however the artifactual reads are not easy to tell apart from normal read duplication due to natural over-sequencing of highly expressed genes. Especially when working with little input material or single cells assessing the fraction of duplicate reads is an important quality control step for NGS data sets. Up to now there are only tools to calculate the global duplication rates that do not take into account the effect of gene expression levels which leaves them of limited use for RNA-Seq data.

**Results:** Here we present the tool dupRadar, which provides an easy means to distinguish artefactual from natural duplicate reads in RNA-Seq data. dupRadar assesses the fraction of duplicate reads per gene dependent on the expression level. Apart from the Bioconductor package dupRadar we provide shell scripts for easy integration into processing pipelines.

**Conclusions:** The Bioconductor package dupRadar offers straight-forward methods to assess RNA-Seq datasets for quality issues with PCR duplicates. It is aimed towards simple integration into standard analysis pipelines as a default QC metric that is especially useful for low-input and single cell RNA-Seq data sets.

## Background

### Sources of duplicate reads in Next-Generation sequencing

Next Generation Sequencing has become a standard assay for many questions in molecular biology. It involves the preparation of sequencing libraries out of fragments of DNA or RNA molecules and sequencing adapters, PCR amplification and sequencing. The calculation of the fraction of duplicate reads has become a standard step for quality control in NGS experiments, as high duplication rates can hint towards problems in different steps of the NGS library preparation process. In particular, the variety of molecules that can be seen after sequencing correlates with minute amounts of input material (“molecular bottleneck”) or too many PCR cycles. This can lead to low library complexity. Furthermore overloading of a sequencing flow cell may result in optical duplicates or problems with reagents can lead to elevated duplication rates. Duplicate reads can also be caused by a combination of complex genomic loci and insufficient read length or even issues with the reference genome.

In RNA-Seq however it is common to have high overall fractions of duplicate reads not due to technical artifacts. This is known and discussed in the community (e.g. [1]) but is still sometimes misunderstood [2]. Often the top 5% of expressed genes take up more than 50% of all reads in a common RNA-Seq dataset [3]. Read counts for highly expressed genes easily surpass the threshold of 1 read per bp of the exon model, at which read duplication is inevitable. Due to a number of biases in the process of RNA-Seq [4] read duplication in RNA-Seq starts even below the 1 read per bp threshold. In RNA-Seq duplication originating from technical artifacts such as described before are confounded with natural read duplication due to highly expressed genes, hence overall duplication rate is not a suitable measure for quality control purposes.

### Effects and treatment of PCR duplicates in RNA-Seq data

In assays involving genomic DNA (e.g. resequencing, ChIP-Seq) reads marked as duplicates with tools such as the established picard [5], or the more recent bamUtils dedup [6] and biobambam [7] are commonly removed before further analysing the data. In RNA-Seq studies with the aim to quantify expression however this is not an option, as highly expressed genes will be affected much stronger by a censoring approach than lowly expressed genes. On the other hand the analysis of RNA-Seq data sets with large amounts of technical duplicates can result in strongly differing analysis results due to the distortion of the relation of expression level and read counts and should not be ignored. Tools such as eXpress [8] attempt to tackle related problems by smoothing the read coverage. However this approach is not applicable to situations in which systematic over-estimation of read counts on a large fraction of genes exists.

### Detection of duplicate reads in Next-Generation sequencing

Currently there are many tools available that address the overall duplication rates or read frequencies of NGS data sets [5,9–14]. Commonly, the non-systematic detection of PCR artefacts in RNA-Seq analysis relies on the visual inspection in a genome browser, where problematic data sets show typical stacked reads in loci with low and medium expression.

Here we present dupRadar, a tool to systematically detect anomalous duplication rate profiles and simplify the task of identification of data sets that require further in-depth assessment.

## Implementation

dupRadar relates the duplication rate and length normalized read counts of every gene to model the dependency of this two variables. It requires a BAM file with mapped and duplicate marked reads, and a gene model in GTF format. Internally dupRadar calls the featureCounts function from the RSubread package [15] several times, to count all and the duplicate marked reads per genes, both uniquely as well as multi-mapping reads. Furthermore dupRadar calculates the per gene duplication rate and reads per kilobase (RPK) as a proxy for relative gene expression. The resulting calculations are stored in a data frame which can be directly passed on to different visualization functions, which show the dependence of the duplication rate on gene expression. Besides fitting a logistic model to the dependency between duplication rate and RPK, dupRadar estimates the baseline duplication rate for lowly expressed genes which can be used as an indicator for general problems inside a data set.

Additionally, the data frame can be used for further processing of the data in standard read count based differential gene expression tools [16–18], or for other purposes such as the detection of genes that are exclusively covered by multi-mapping reads.

To enable interpretation of the dependency of duplication rate and gene expression, dupRadar currently includes various visualization functions. Beyond that the vignette of the Bioconductor package contains examples for customised plots using dupRadar. For the sake of usability, it includes wrappers for some common tools for duplicate marking in order to streamline the processing of the data sets.

## Results and Discussion

Recently, RNA-Seq protocols were improved considerably, leading to less technical duplicates and the linked issues. Still in our experience possible problems are worth to be checked for by default, especially if protocols are pushed to or beyond their boundaries or more recent low-input or single cell RNA-Seq protocols are used.

To demonstrate the usage of dupRadar we apply a typical work flow for selected single read RNA-Seq data sets from the study of Marinov et al. [19] ranging from single cells to cell pools to bulk RNA data. We map reads using STAR [20] and mark duplicate reads using BamUtils dedup [6]. Together with the human reference gene annotation GTF included in the iGenomes collection for the UCSC hg19 build [21], we use the resulting bam files as input for dupRadar’s duplication rate calculation function. As an example Tab. 1 contains the entries from a sample of 10 genes out of the full set for the library 13276 (SRR764800).

**Tab. 1:** Example values for a sample of 10 genes from the library 13276. Some columns were omitted due to space constraints; refer to Suppl. table 1 for the complete table.

| ID | geneLength | allCounts | filteredCounts | dupRate | dupsPerId | RPK | RPKM |
| --- | --- | --- | --- | --- | --- | --- | --- |
| LOC100288069 | 1371 | 17 | 15 | 0.12 | 2 | 12.40 | 0.60 |
| LINC00115 | 1317 | 28 | 28 | 0.00 | 0 | 21.26 | 1.03 |
| LOC643837 | 9233 | 281 | 246 | 0.12 | 35 | 30.43 | 1.47 |
| FAM41C | 1706 | 1 | 1 | 0.00 | 0 | 0.59 | 0.03 |
| LOC100130417 | 496 | 0 | 0 | NA | 0 | 0.00 | 0.00 |
| SAMD11 | 2554 | 0 | 0 | NA | 0 | 0.00 | 0.00 |
| NOC2L | 2800 | 329 | 273 | 0.17 | 56 | 117.50 | 5.67 |
| KLHL17 | 2564 | 2 | 2 | 0.00 | 0 | 0.78 | 0.04 |
| ISG15 | 666 | 590 | 271 | 0.54 | 319 | 885.89 | 42.78 |
| AGRN | 7326 | 3 | 3 | 0.00 | 0 | 0.41 | 0.02 |

Based on the duplication rates, we generate the main visualizations of dupRadar in Fig. 1. The effects of over-sequencing libraries of limited complexity in cases of little input material as well as an example for a bulk RNA-Seq dataset without any traces of PCR duplicates. The given plots indicate the duplication rate in relation to the gene expression. Usual single read RNA-Seq experiments at common read depths are expected to show low duplication rates for lowly expressed genes in the bottom left of the plot, with the duplication rate rising as the expression level approaches the 1 read/bp boundary. Beyond this threshold genes are covered almost completely with reads marked as duplicates due to their high expression levels (e.g. Fig. 1C). Data sets based on lower amount of input material show the effects of limited complexity of the library, resulting in higher duplication rates already at lowly expressed genes, leading to the majority of data points being shifted upwards to higher duplication rates also for lowly expressed genes (e.g. Fig 1D). Similar situations can be observed for data sets with actual PCR artifacts.

**Fig. 1.**
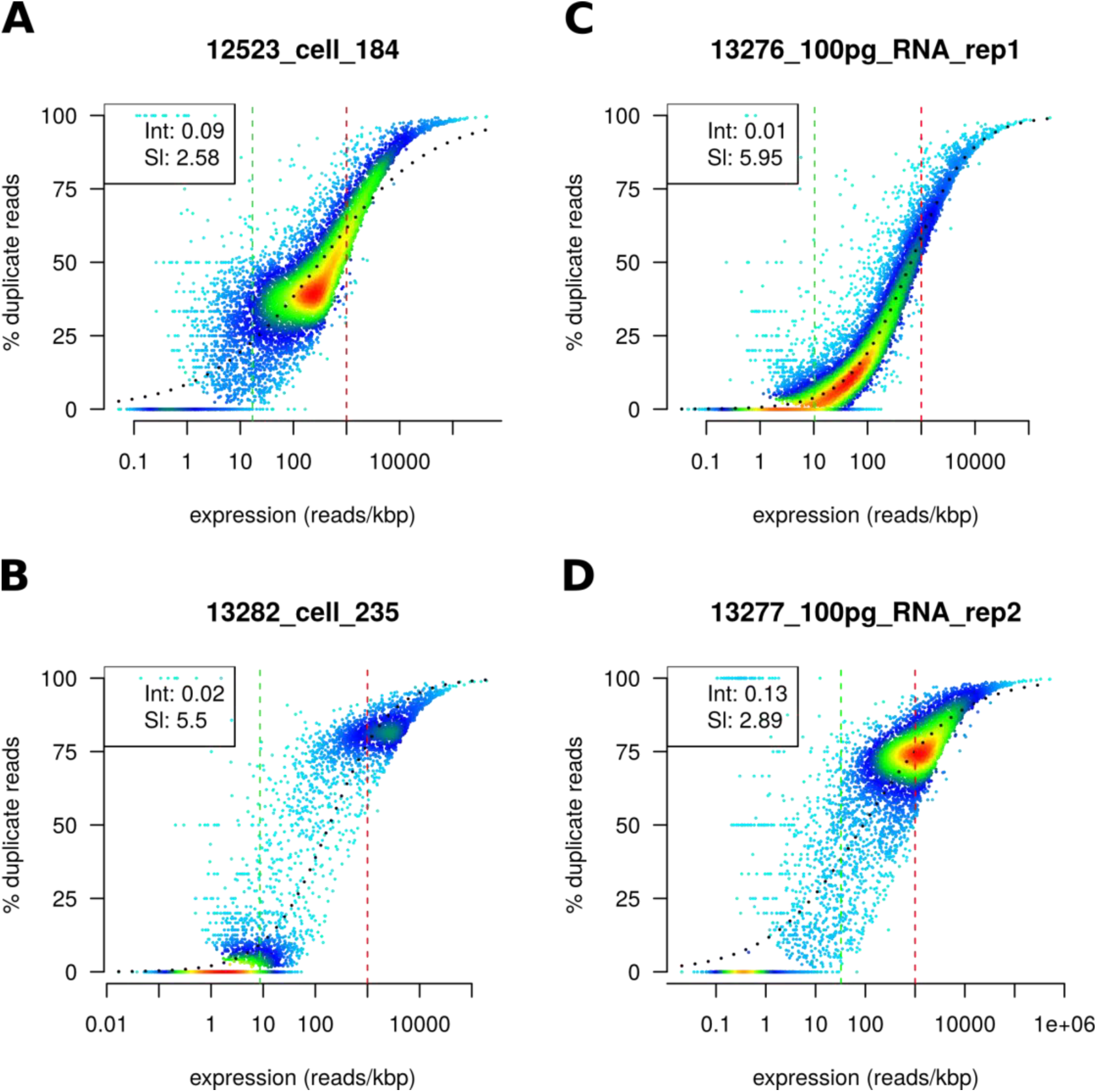
Several RNA-seq datasets from Marinov et al.[19]. Legends shows the intercept and slope of a fitted logit model. **A** Single cell experiment with relatively low duplication rates and most of the genes detected. **B** Single cell experiment with most of the genes undetected and high duplication rate on the detected ones. **C** RNA-seq experiment pushing the protocol to only 100pg of input material, with low duplication rates and relatively good identification of genes. **D** same RNA-seq experiment, showing over-sequencing due to higher sequencing depth of the library.

Although paired-end libraries facilitate the distinction between duplicates due to adding the fragment length as an extra variable to distinguish molecules, still the problem is not completely solved. The recent introduction of unique molecular identifiers (UMI) during library preparation, allows for exact distinction of technical and biological duplicates and therefore also the removal of technical duplicates [22], which alleviates the described problem on the side of experimental procedures.

## Conclusions

The Bioconductor package dupRadar offers straight-forward methods to assess RNA-Seq datasets for problems with duplicate reads and is aimed towards simple integration into standard analysis pipelines as a default QC metric.

While dupRadar serves as a diagnostics method for PCR duplicates, we regard the issue of correction for these artefacts as yet unsolved, with a potential to extend dupRadar with correction functions. Currently we advise colleagues to treat with caution RNA-seq data strongly affected by technical duplicates and repeat library preparation and sequencing if possible.

Similar effects comparable to over-sequencing of highly expressed genes are implicated for certain types of enrichment-based assays (e.g. ChIP-Seq of a specific transcription factor with high read-depths). Suitability of dupRadar in this area remains to be explored.

## Availability

- Project name: dupRadar
- Project home page: http://bioconductor.org/packages/dupRadar/
- Operating system(s): Linux; MacOS
- Programming language: R >= 3.2
- Other requirements: Bioconductor>= 3.2
- License: GNU GPL
- Any restrictions to use by non-academics: None

## List of abbreviations

RPK: reads per kilobase
PCR: polymerase chain reaction
UMI: unique molecular identifiers
QC: quality control
ChIP: chromatin immunoprecipitation
bp: base pair
NGS: next-generation sequencing

## Authors' contributions

SS and HK conceived of the project. SS, HK designed the software. SS, DS and HK implemented the software. SS and HK tested the software. SS and HK drafted the manuscript. All authors read and approved the final manuscript.

## Acknowledgements

We thank the members of the Core Facilities at the Institute for Molecular Biology, especially Joern Toedling, Nastasja Kreim, Anke Busch, Oliver Drechsler and Emil Karaulanov, as well as Germán Leparc from the Computational Biology Group at Boehringer Ingelheim for discussion, input and proof-reading.

